# Rationally Designed, ACE2 Mimetic Binder to the SARS Cov-2 Associated Spike Protein for COVID-19 Therapeutics and Beyond

**DOI:** 10.1101/2025.06.05.658001

**Authors:** Michael H. Peters, Xin Yan Pei

## Abstract

We have developed a 23 residue, optimized helical peptide biomimetic from the SARS-CoV-2 Spike protein binding partner ACE2 that demonstrated a Therapeutic Index (TI) of >∼20 in authentic viral cell challenge assays, including WT and Omicron strains. The therapeutic is an optimized “peptide decoy” based on the virus’s human cell target ACE2 and, as such, may have more general applicability across coronavirus family members that use ACE2 for cellular entry. We experimentally verify a comprehensive, *rational* optimization strategy of the peptide through improved binding, helical content, and solubility from its native sequence. These techniques may have general applicability for helical peptide optimization for other therapeutic targets as well. Importantly, techniques also exist for protecting helical peptides in vivo for improved delivery. Peptides are readily modifiable with single residue substitutions for quick response to mutated targets, and they typically have relatively low toxicity and ease of manufacturing, making peptides extremely attractive as biological therapeutics against viral pathogens. The general concept of using peptide decoys across other viral human cell targets is also discussed.

**Author Summary:** Although the COVID-19 pandemic has ended, SARS-CoV-2 virus still causes hundreds of deaths each week world-wide. The virus uses a human cell receptor called ACE2 to latch on and enter cells. Here, the small critical attachment segment of ACE2 is developed as a “decoy” molecule thereby protecting host cells and significantly reducing viral replication, which is the role of a therapeutic agent. The effective segment in this case is called a helical peptide and it was optimized for better binding, solubility, and stability using rational methods based on our understanding of this class of molecule.

Moreover, future viruses from this family of coronaviruses may likely use ACE2 as their host cell receptor, as recently demonstrated in the Middle Eastern Respiratory Syndrome Virus of bats and, therefore, the ACE2 decoy therapeutic may have future applications as well. Development of a broader range of decoy molecules for different viruses using different human cell receptors may represent a forward-thinking approach to prepare for future pandemics and outbreaks.

## Introduction

SARS-CoV-2 is a β coronavirus responsible for over 578 million infections and over 6.9 million deaths world-wide. Although no longer a pandemic classification, its current circulating variants continue to pose threats through potential resistance to existing therpaeutics and vaccines. β coronaviruses represent one (B) of four genera (A,B,C,and D) of RNA positive-sense viruses in the Nidovirales order ^1,2^. SARS-CoV-2 is the latest in human viral outbreaks of this genera being preceded by the Middle Eastern Respiratory Coronavirus (MERS-CoV) and the SARS-CoV outbreak of 2002.

Quantitatively, each SARS-CoV-2 virion carries approximately 100 Spike proteins per virion^3^. The prefusion Spike protein (S) is a large trimeric protein where each protomer may be in a so-called Up state or Down state, depending on the configuration of its receptor binding domain (RBD). In the “Up-state” the (prefusion) protein is able to efficiently bind to human epithelial cells expressing Angiotensin Converting Enzyme 2 (ACE2) on their surface and infect via a transformation to its fusion state. These cells include Type I and II pneumocytes; alveolar macrophage; nasal mucosal cells; and a large range of vascular epithelial cells throughout the body (kidney, heart, etc.).^4^ In addition, interaction of the Spike protein with TMPRSS2 and possibly other receptors may be another important route to viral entry^5^. However, ACE2 has been shown to play *the critical role* for viral entry^5,6^.

In addition, it is well-known that this family of β coronaviruses has the capability of recombination within animal hosts, leading to potential zoonotic transmission of recombinant viruses^7,8^. For example, it has been recently discovered in bats that a variant of MERS-CoV uses the ACE2 receptor, like SARS-CoV-1 and SARS-CoV-2, to infect human cells^9,10^. Other human coronaviruses also use the ACE2 receptor for cellular entry^11^. The disturbing situation is the broad range of potential viral threats from this family of viruses as a result of the “soup” of variations and recombination’s possible in the environment, and the potential subsequent transmission to humans.

The concept of pan-coronavirus vaccines and therapeutics looks to exploit the common features of viruses that can be used as conserved molecular targets across viral variations. It is clear that families of viruses use specific human cell receptors for their entry, and the development of cell receptor molecular decoys represents one possible path to pan-coronavirus therapeutics. Table 1 summarizes some of the viruses and their known human cell receptor host molecules. Human cell receptors have been used to develop decoy therapeutics, such as sialic acid mimetics for Influenza A^12^, ACE2 for SARS, and others such as ICAM-1^13^. Human cell receptors typically have key binding sites that may be conserved across viral mutants, variations, and recombination events.

**Table 1.** Examples of Human Cell Receptors for Viral Entry.

| **Virus Name** | **Human Cell Receptors** | **Central References** |
| --- | --- | --- |
| **SARS-CoV-2** | ACE2 (Angiotensin-Converting Enzyme 2) | Hoffmann et al., 2020<sup>14</sup>;<br>Wrapp et al.<sup>15</sup>, Walls et al.,<br>2020<sup>16</sup> |
| **Influenza A** | Sialic acid receptors (α2,6 and α2,3 linkages) | Rogers et al., 1983<sup>17</sup>;<br>Matrosovich et al., 2004<sup>18</sup> |
| **Rhinovirus** | ICAM-1 (Intercellular Adhesion Molecule 1),<br>LDLR (Low-Density Lipoprotein Receptor) | Greve et al., 1989<sup>19</sup>; Hofer et al., 1994<sup>20</sup> |
| **Measles virus** | CD46, SLAM (Signaling Lymphocyte Activation Molecule), Nectin-4 | Tatsuo et al., 2000<sup>21</sup>;<br>Mühlebach et al., 2011 |
| **Respiratory Syncytial Virus (RSV)** | Nucleolin, CX3CR1 (Fractalkine receptor) | Tayyari et al., 2011<sup>22</sup>; Johnson et al., 2015<sup>23</sup> |
| **Human Metapneumovirus (HMPV)** | Heparan sulfate, Proteoglycans | Chang et al., 2006<sup>24</sup>; Cox and Williams., 2013<sup>25</sup> |
| **Adenovirus** | CAR (Coxsackievirus and Adenovirus Receptor), Integrins (αvβ3, αvβ5) | Wickham et al., 1993<sup>26</sup> |

| Virus Name | Human Cell Receptors | Central References |
| --- | --- | --- |
| **Varicella-Zoster Virus (VZV)** | Insulin-Degrading Enzyme (IDE) | Zerboni et al., 2014<sup>27</sup> |
| **Human Coronavirus OC43** | HLA Class I Molecule, Sialic acid receptors | Hulswit et al., 2019<sup>28</sup> |
| **MERS-CoV** | DPP4 (Dipeptidyl Peptidase 4) | Raj et al., 2013<sup>29</sup> |

In this study, the key binding segment of human ACE2 is exploited to develop an optimized peptide that efficiently mimics the key binding segment of ACE2 with the receptor binding domain (RBD) of the spike protein of SARS-CoV-2. Although ACE2 derived peptides have been proposed and studied as SARS-CoV-2 therapeutics^30,31^, these studies are often based on the use of large compound libraries and random structure variations (high throughput screening). Note that the native α1 helix was studied some time ago as a potential therapeutic to SARS-CoV-1^32^. Small segments of the native peptide (∼6 residues in length) have also been studied as SARS-CoV-2 viral inhibitors, such as E37-Q42 hACE2 segment^33^. Here a rational approach for optimizing the native peptide is taken based on understanding of binding physics, helical structure/stability, and peptide solubility. These methods may also be transferrable to other targets and biological therapeutics as an alternative to large compound libraries and high throughput analyses, which can be costly and time-consuming.

## Results

In order to generate optimized peptide mimetics to the RBD of the Spike protein, all-atom, *ab initio* molecular dynamics were initially carried out over a 0.5 microsecond total time based on the published structure of the RBD of SARS-CoV-2 Spike protein (S) with ACE2^34^ (PDB ID 6M17; B and E chains), as described previously^35^. The average conformation of the binding complex near the end of the simulation is used to map the dominant energetic atom-atom interactions between ACE2 and the RBD as shown in Fig. 1. The dominant interactions in this case are the partial charge interactions between ACE2 and RBD residues among both backbone and side chain atoms of the amino acid residues, as shown in Fig. 2. There are two distinct binding domains to ACE2 : residues 22 -41 (alpha helix 1) and residues 322-357 (loop domain).

**Figure 1.**
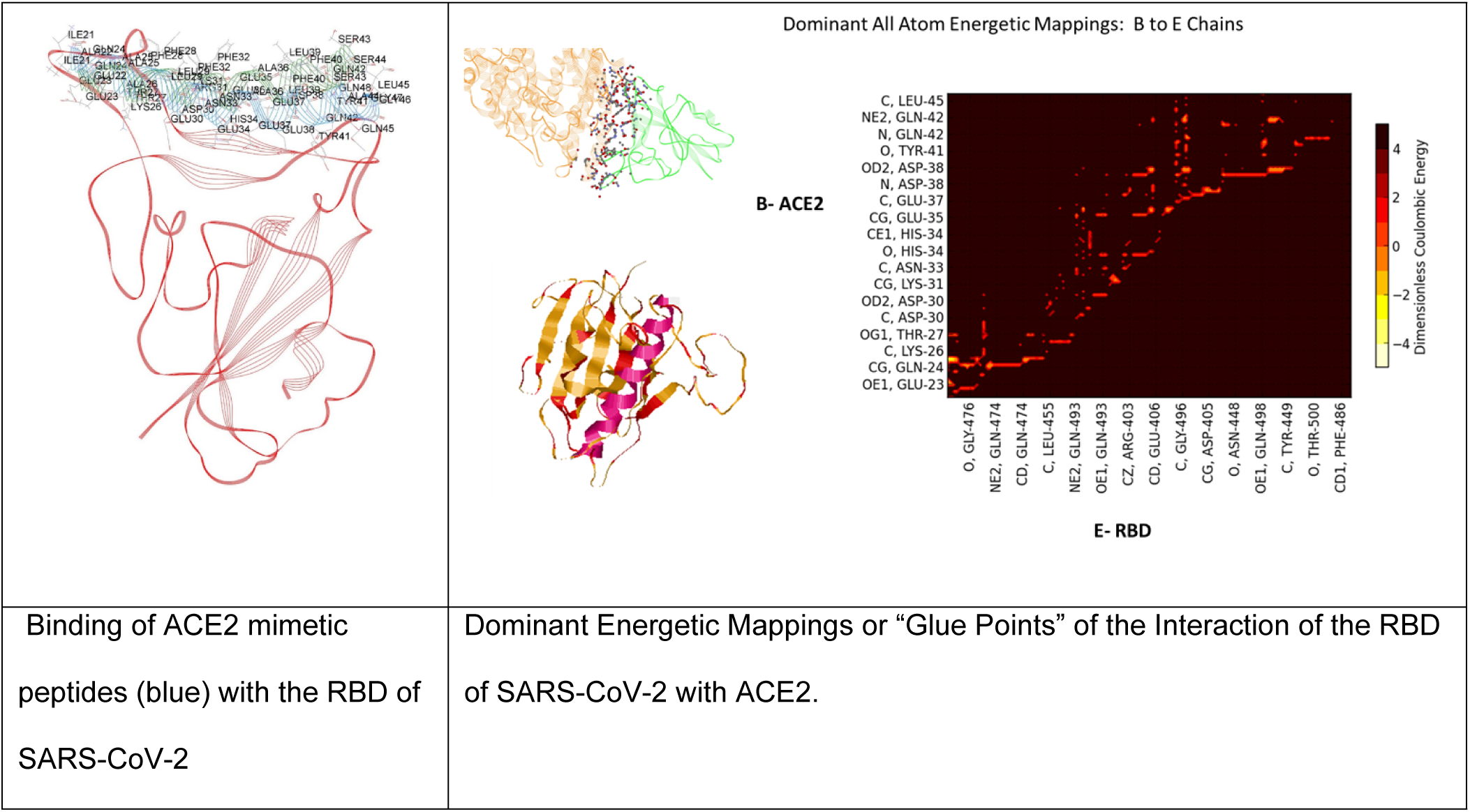
Dominant energetic mappings for the development of peptide biomimetics from ACE2^35^. Complete quantitative information is given in Supplementary Data S1.

**Figure 2.**
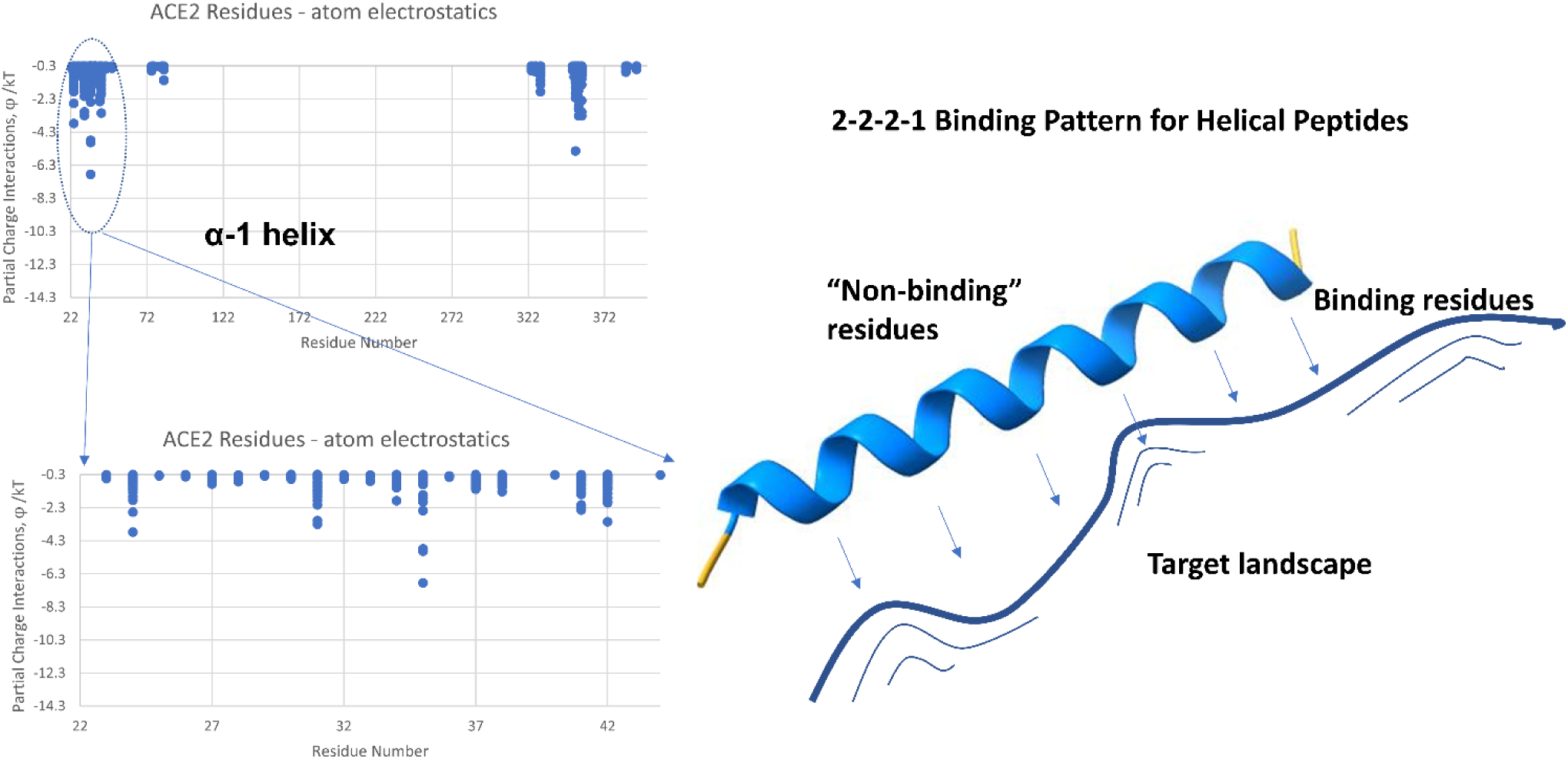
Binding of SARS-CoV-2 to human ACE2 showing the two distinct binding regions: α1 helix (residues 23-42) and loop domains (324-357). Binding energies are normalized by kT, where k is Boltzmann’s constant and T is absolute temperature.

Specific energetic contact points (atom-atom interactions) for helical binding typically form a distinct pattern of 1 residue “on”, 2 residues “off”, 2 on, and 2 off. The pattern repeats approximately every 7 residues owing to the helical geometric requirement of 3.4 residues per turn, as shown in Fig. 2

Unfortunately, the α1 helix of ACE2 unfolds in isolation from its parent protein and no longer maintains its helical structure, and is a weak binder to spike RBD in isolation. There are also solubility issues owing to the relatively large number of negatively charged groups. In the native state, the so-called non-binding residues (with respect to the RBD of Spike) in Fig. 2 strongly interact with the α2 helix of ACE2 helping to stabilize its helical structure. In addition to improving binding to the Spike RBD through rational examination of single residue substitutions for binding residues, non-binding residue substitutions can be used to promote helicity and solubility. Table 2 summarizes the results of the rational strategy to optimize the peptide sequence, Fig. 3 shows a side-by-side computational comparison of the optimized to native helical binding behavior, and Fig. 4 shows a helical wheel comparison.

**Figure 3.**
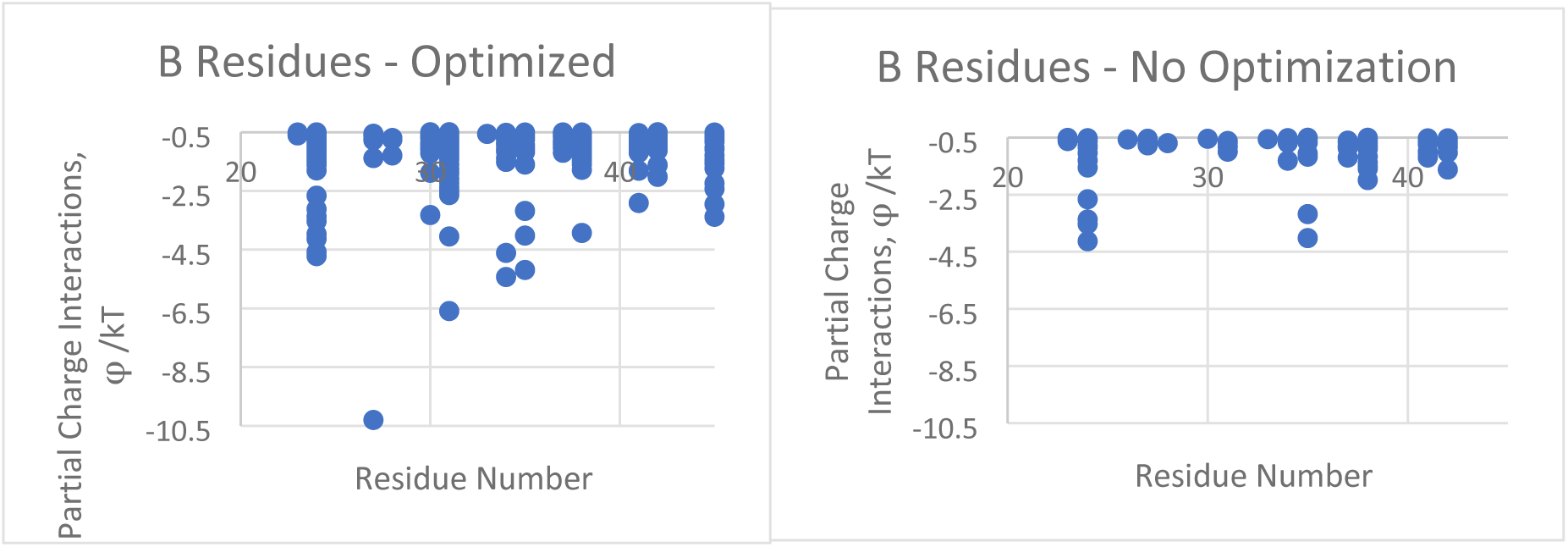
**Right**: Partial charge interactions of the native residue α1 helix segment of hACE2 with the RBD of SARS-CoV-2. **Left:** Significantly improved interactions with Spike RBD after optimization for the isolated, 23-residue peptide (native residues 22 through 44). Some substituted residues are associated with improved helical stability and solubility. Helical residues (<∼30 residues in length) form a unique 7 residue repeat sequence due to the requirement of 3.4 residues per turn which is exploited in the optimization process (Fig. 4).

**Figure 4.**
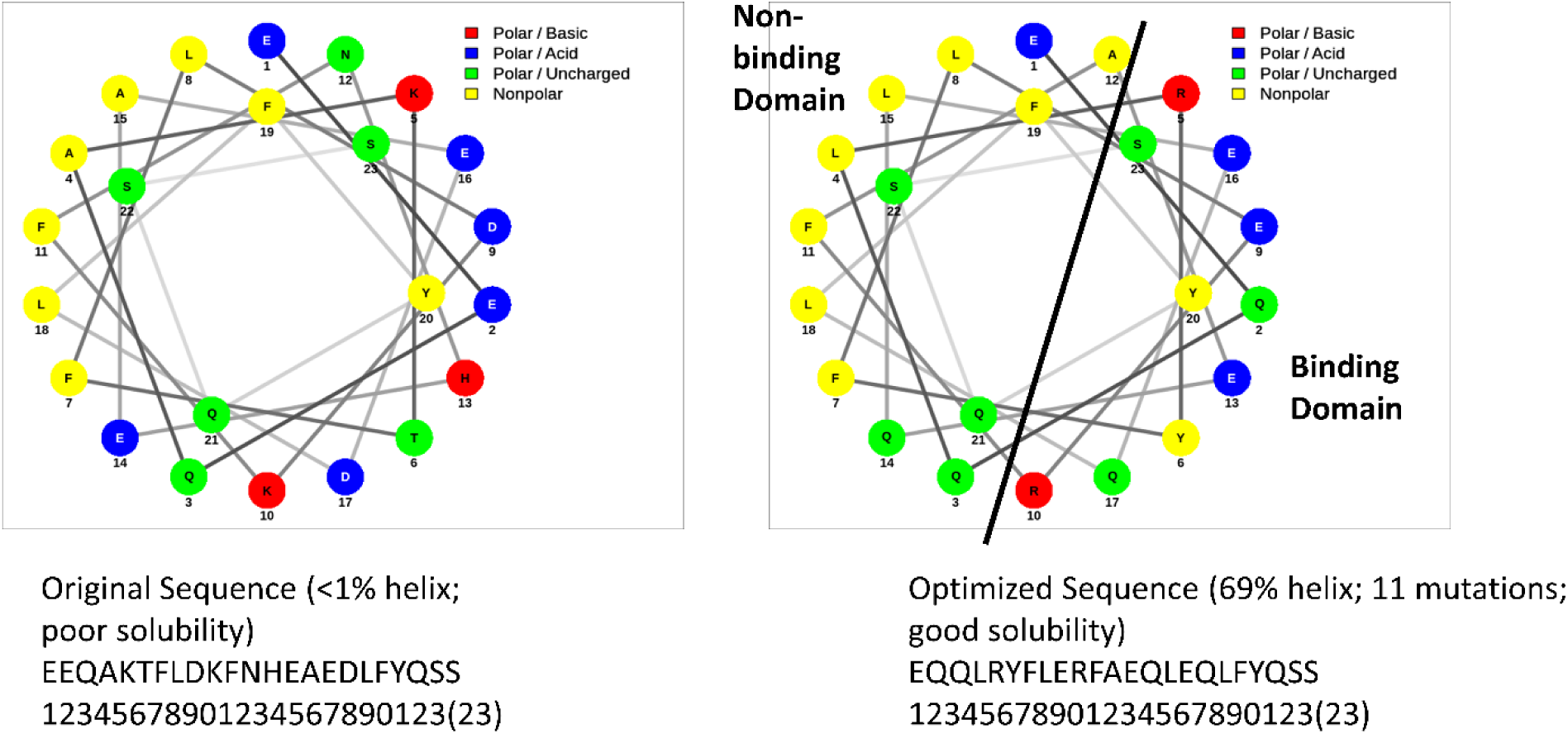
Helical wheel representation of native and optimized sequence. Binding domain versus non-binding domain is shown in the optimized representation. (Helical wheels generated by NetWheels^36^)

**Figure 5.**
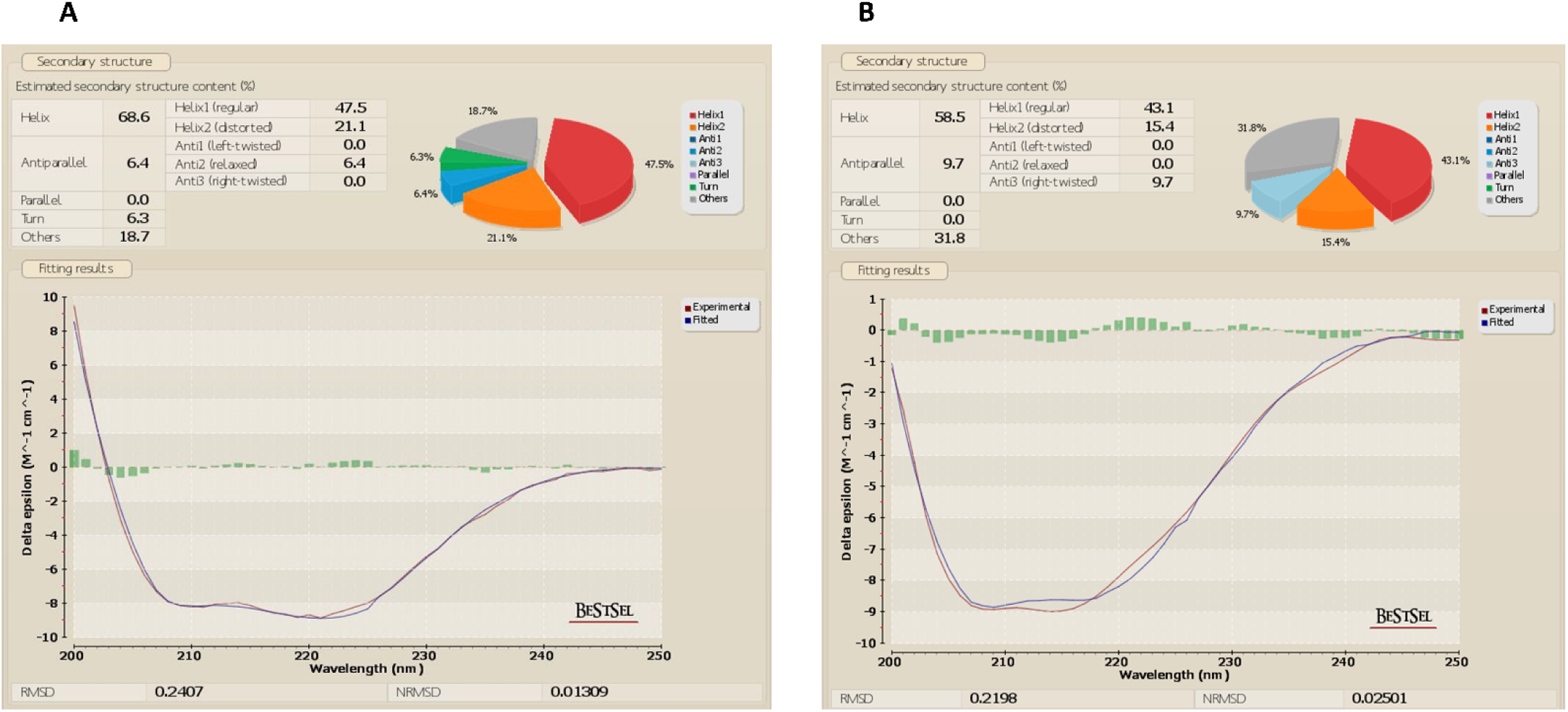
Experimental Circular Dichroism (CD) measurement of A. AK26R peptide. B. A peptide (Graph and analysis from BestSel^40^)

**Table 2.**
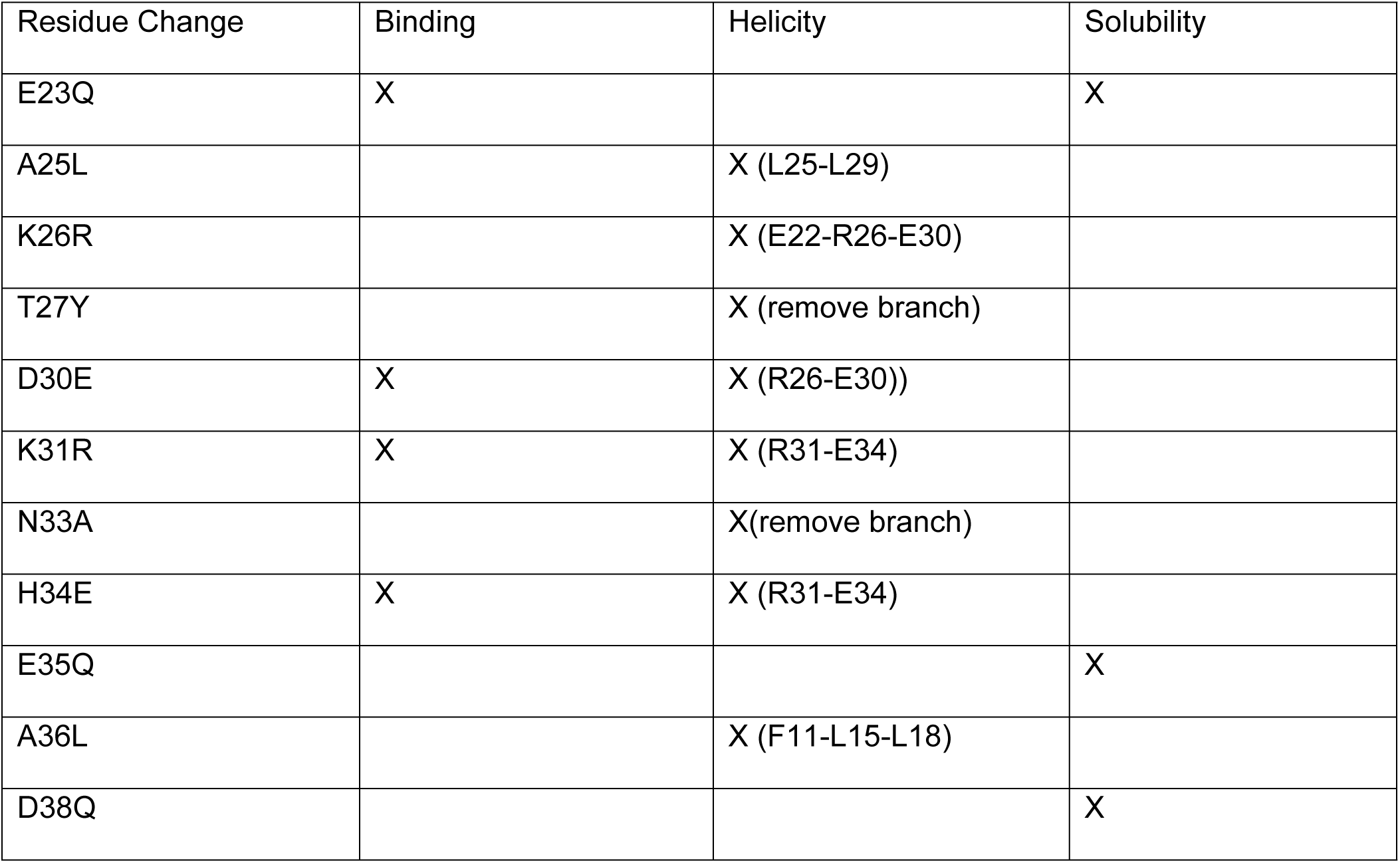
Summary of Single Residue Changes from The Native State.

Helicity strategy follows the study of Facchiano et al.^37^ and Creamer and Rose^38^ Here, E-K at positions (i, i+3) or (i, i+4) (charge “staple”), L-L and L-F at (i,i+3) or (i, i+4) (non-polar “staple”), and branch removal were used for this particular native sequence. Rational, in-silico single residue substitutions to improve helicity were computationally verified using Agadir^39^. Note that N-terminal acetylation and C-terminal amidation were included. It was found here that E-R can be a slightly more effective staple then E-K for this particular sequence perhaps due to the small rotational change allowed for the protonated amine side chain of arginine. Fortunately, the three binding improvements shown in Table 2. also resulted in improved stapling effects in this particular case. Binding improvement strategy involves improving the specific charge or partial charge interactions of the helix with the RBD, such as H34E shown in Table 2, which was strategized through atom-atom interaction mappings and verified computationally (see Methods).

The water solubility, in this case, was addressed through reduction in the total number of charged residues from a -3 native charge to a -2 charge in the optimized helix and, at the same time, keeping the number of polar and charge groups greater than the number of hydrophobic residues (see Fig. 4).

Computationally optimized peptides were then physically synthesized for in-vitro studies. In this case, two peptides with and without K26R were studied (Peptide A and AK26R). In the end, AK26R, experimentally showed slightly better helicity and authentic viral inhibition, but both peptides were effective at Spike RBD binding, dissolution in water, and significantly improved helical structure. Note that the experimental results reported here were obtained in real-time over the course of the pandemic. Because of the frequency of change, studies on all of the different variants of concern (VOC) that appeared during the pandemic are beyond the scope of this work.

Experimental CD measurements of optimized peptides demonstrated excellent helicity (68.6% AK26R; 58.5% A), as shown in Fig. 4. The small overall charge reduction from -3 to -2 had a significant effect on improving solubility. Although not quite fully soluble in nanopure water, the addition of as little as 4% of a 1.0 M solution of NaOH completely dissolved the optimized peptides in water.

Experimental binding studies of optimized peptides were carried out over a range of SARS-Cov-2 variants during the course of the pandemic. This is illustrated in Fig. 6 for a set of variants prior to the appearance of the Omicron variant (see Supplemental S3 for additional SPR studies). Equilibrium dissociation constants, K_D_ =(k_d_/k_a_, dissociation/association kinetic rate constants) demonstrated the effectiveness of the optimization strategy using ACE2 α1-helix, which all variants must use for host cell entry.

**Figure 6.**
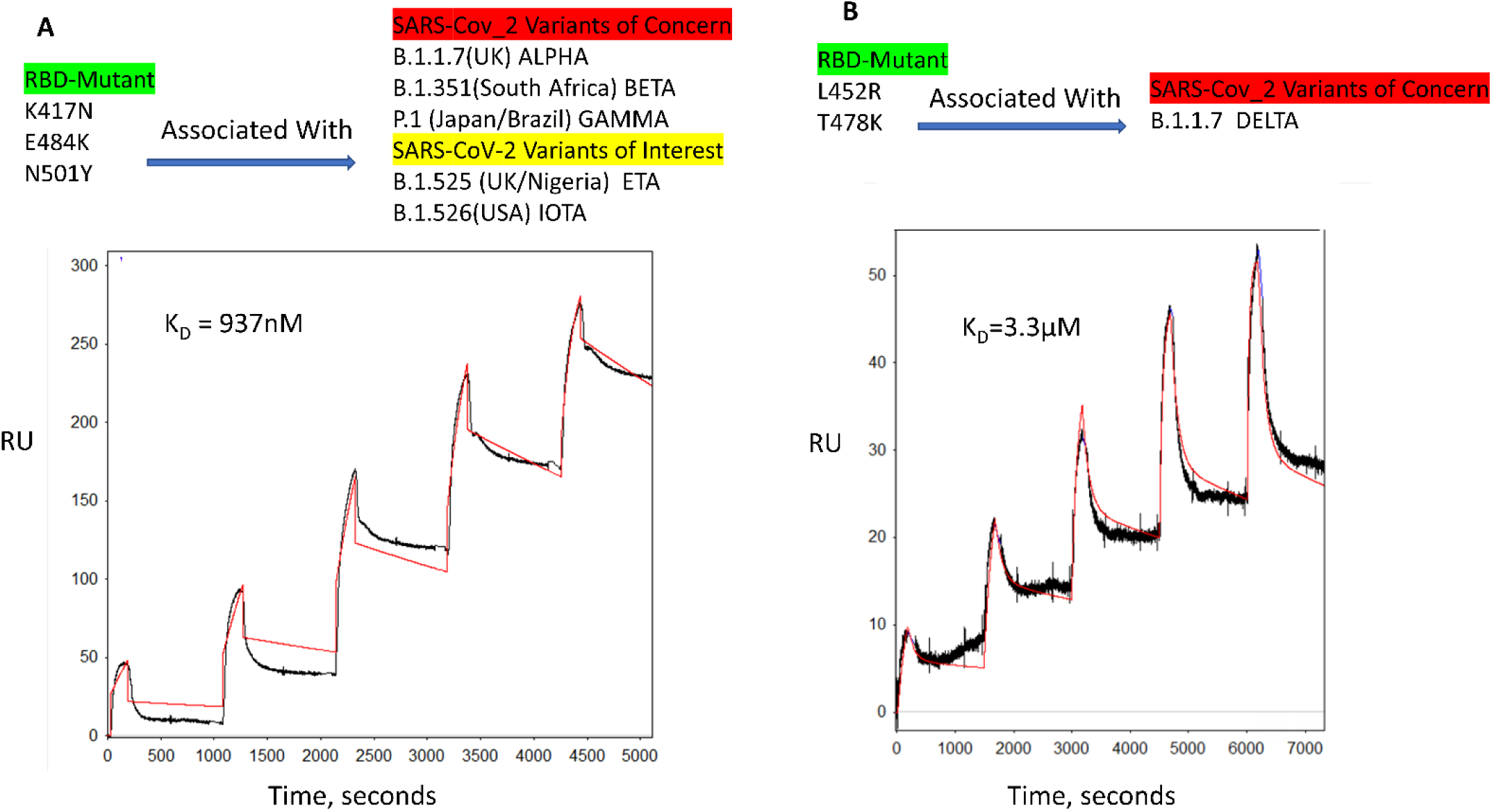
A. Binding of Peptide A to RBD-mutant (K417N, E484K, and N501Y) associated with a number of SARS-CoV-2 variants of concern during the pandemic, as measured by Surface Plasmon Resonance (SPR; see Methods). B. Peptide A binding to the Delta variant as measured during the pandemic. SPR data of hACE2 binding to the Omicron variant as positive control is given in Supplementary S3.

Authentic viral inhibition studies were subsequently carried out during the course of the pandemic in a BSL3 facility with the two lead, optimized peptides against Wuhan WT and what became the dominant circulating variant: the Omicron variant. As shown in Fig. 7, peptide A was effective against the WT

**Figure 7.**
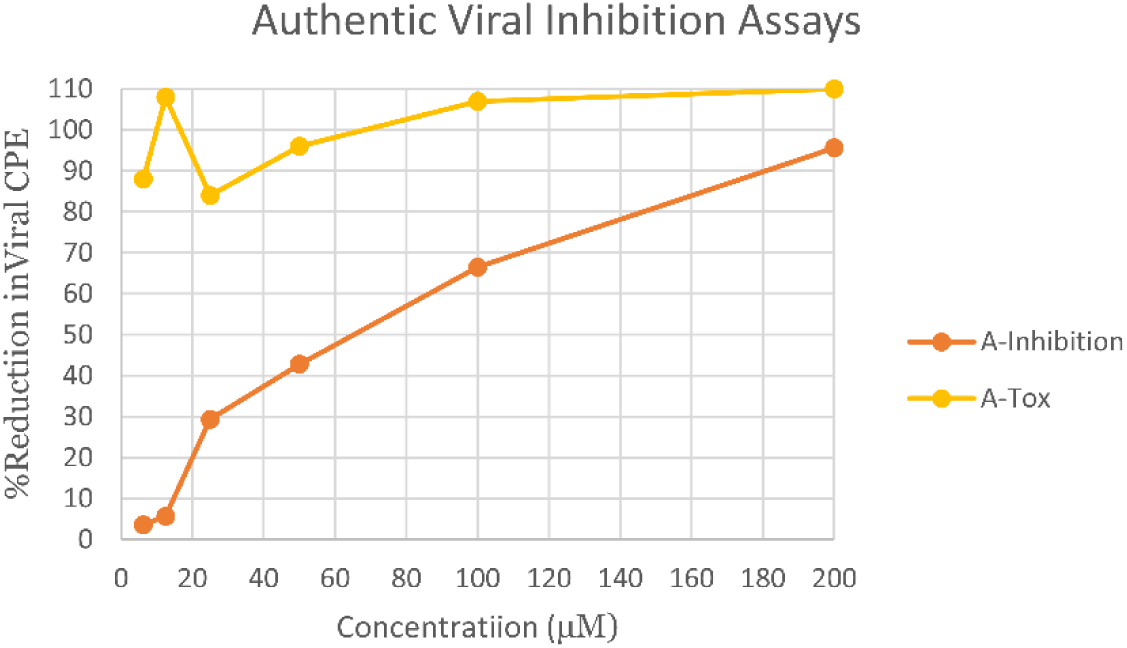
Authentic Viral Cell Challenge Assays as measured during the pandemic. Peptide A was tested against Wuhan WT (USA_WA1/2020) (Therapeutic Index >∼4) (Red). Peptide AK26R was tested against Omicron BA.2. (Therapeutic Index >∼36) (Supplemental S4). No toxicity to cells by either peptide in the absence of virus, was observed. (Yellow). The coefficient of variation (COV) across peptide A inhibition studies averaged ∼60%, whereas AK26R inhibition studies averaged ∼40%; all toxicity values reported were ∼10% COV (Supplemental S4).

Wuhan SARS-CoV-2 strain (TI>∼4). Peptide AK26R was also effective against Omicron BA.2 with a Therapeutic Index >∼36 (Supplemental S4). Again, these were the experimental activities carried out in real-time during the course of the pandemic. Optimized peptides have also been exposed to other cell lines, such as A549 human lung cells (Supplemental S2), with no cellular toxicity out to millimolar levels of peptide concentration leading to much larger TI’s.

Repeating of authentic viral inhibition tests are currently planned against circulating Omicron sub-lineages and for testing effectiveness in pre-clinical animal models. Note that authentic viral inhibition tests were also done in parallel using the known IV therapeutic, Remdesivir, as a comparison at the time of these studies (Supplemental S4). Because all variants use the ACE2 receptor, it is expected that the optimized peptides would continue to remain effective across future strains and possibly other emerging viruses that use the human ACE2 receptor, as discussed more fully below.

## Discussion

Here we have demonstrated a rational approach to the optimization of a helical biomimetic of the ACE2 receptor domain for the possible use as a therapeutic agent in the treatment of COVID-19 and other viruses that use human ACE2 for cellular entry. The importance of seeking small peptide therapeutics or prophylactics is centered on ease and rapidity of world-wide manufacturing, ease of administration, such as intranasal for airborne viruses, and the peptides demonstration of extremely low toxicities. The optimization method begins with large scale, *ab initio* molecular dynamic computations and the mapping of specific atom-atom interactions between the ACE2 receptor and spike RBD that are responsible for the binding behavior. Helical peptides have a binding domain and a non-binding domain with a specific pattern owing to the regular geometry of 3.4 residues per turn. The non-binding residues may be changed to promote helicity according to known residue interactions that promote helical formation. Solubility issues can also be addressed via residue substitutions in the non-binding domain, as demonstrated here. Since all variants use the ACE2 receptor for entry, the approach is demonstrated to withstand spike RBD variants. Although not guaranteed, the α1 segment of ACE2 is the exposed native binding segment of this receptor and may be likely to remain so for future variants or viral recombination.

Viruses recombine through genetic exchange mechanisms that allows mixing and shuffling of genetic material, leading to the emergence of new viral strains^7,8,41^. Recombination can occur between different viral strains of the same type or differing viral types; in either case with the same host cell. The primary modes of recombination in RNA viruses include template shifting in non-segmented genomes and reassortment in segmented genomes^8^. It is noted that SARS-Cov-2 is a non-segmented genome, whereas Influenza A is an 8-segmented genome. In non-segmented RNA viruses, recombination primarily occurs through a mechanism called "copy choice recombination," where the RNA polymerase, during replication, switches from one RNA template to another^8^. Reassortment involves mixing of the genome segments between two different strains or two different types of viruses during host cell viral reproduction^7,41^. Recombination plays a crucial role in viral evolution, immune evasion, and cross-species transmission, contributing to the emergence of novel and potentially more virulent strains.

Although the vast majority of viral recombination effects are deleterious in nature, it is reasonable that non-deleterious recombination would likely conserve human cell surface host receptors in order to be effective at human transmission. Additionally, viral surface protein mutations alone could lead to zoonotic transmission to human cell receptors, for example, “Bird Flu” or H5N1 influenza is currently one mutation away from switching from its current avian sialic acid α2-3 receptor to a human α2-6 sialic acid receptor^42^ (Table 1); again, giving credence to using forms of human cell receptors as therapeutic or prophylactic agents.

Coronaviruses and Influenza A viruses appear to present the highest likelihood for recombination events that could lead to pandemics due to their genetic recombination mechanisms, frequent zoonotic events, and capacity for human-to-human transmission^43^. Other viruses, like Paramyxoviruses (Nipah), Orthomyxoviruses (Thogotovirus), and Flaviviruses (Dengue and Zika), also have the ability to infect humans and co-circulate in high-risk areas where recombination may produce new pathogenic variants^44^. Development of cell receptor decoys may represent a forward-looking and robust approach to viral therapeutics beyond coronaviruses, for example, as recently demonstrated for the Respiratory Syncytial Virus (RSV) human cell receptor, ICAM-1^13^. Note that ICAM-1 is also used by the Rhinovirus (Table 1).

From a practical bent, it is noted that peptides less than 30 residues in length can be manufactured relatively quickly for clinical use in large scale peptide synthesizers (non-biologic). Those synthesizers are available world-wide, which affords world-wide therapeutic production without supply chain issues, which is of great importance during world-wide viral outbreaks. Single residue substitutions of decoy peptides can also be readily invoked to improve binding against any emerging variants. Specific in-vivo therapeutic delivery methods are not addressed here, but intranasal, inhalation, oral, and subcutaneous pathways are well-known, for example, as in the current popular GLP-1 agonist drugs (Victoza, Wegovy, etc.)^45^. GLP-1 (Glucagon Like Peptide 1) is a small helical peptide that is subcutaneously administered and protected in-vivo by the addition of a fatty acid group. Thus, pharmacokinetic issues with peptides as therapeutics have also been addressed over the years. Furthermore, developing several potential weapons against possible future emerging viral pathogens would appear prudent given the variability in patient response to any one particular intervention.

## Materials and Methods

### Peptides and Proteins

Peptides were synthesized to 95+% purity by GenScript (NJ). SARS-CoV-2 spike proteins were purchased through ACRO Biosystems (DE).

### SPR Kinetic Binding Studies

Kinetic binding assays were performed using a Reichart Surface Plasmon Resonance (SPR) instrument (Reichart, NY) according to the manufacturer’s guidelines using a Streptavidin (SA) immobilization chip. RBD WT and VOCs are synthesized and biotinylated at the C-term end (ACROS Biosystems) to avoid any blocking of the binding domains. Briefly, the RBD WT or RBD VOCs were immobilized to the SA chip surface at a concentration of 25 μg/ml and flow rate of 10 μl/min in running buffer. For peptide kinetic binding measurements to RBD, peptide at various concentrations in running buffer were injected over immobilized RBD at a flow rate of 25 μl/min for 3 min with dissociation time of 20 mins. Because of the slow off-rate, strong peptide binding, and despite repeated attempts using a suite of suggested regeneration buffers, the “CLAMP” method had to be employed for all runs due to the difficulty of removing adsorbed peptide without damaging the RBD protein^46^. All data was reference subtracted and fitted to 1:1 Langmuir binding model in “CLAMP” to obtain the association rate constant (k_a_), dissociation rate constant (k_d_) and the equilibrium dissociation rate constant (K_D_).

### Circular Dichorism (CD) and Peptide Helical Content

Peptide helical content was experimentally measured using a Jasco J-1500 CD spectrometer according to the manufacturer’s guidelines. Briefly, samples were measured at concentration of 0.1mg/ml in a 1 mm path length quartz glass cuvette.

Background (dilution buffer) readings are reference subtracted and helical content is determined in the wavelength range of 175 to 300 nm using the BESTSEL software^40^. Computationally, helical peptide content was estimated from the AGADIR software which has been shown to compare well to experimental helical content (2%±6%)^47^.

### Authentic Viral Inhibition Studies

Authentic viral inhibition studies were carried out in BSL3 facilities (ImQuest Biosciences, Inc.). Optimized peptides were tested for antiviral activity against SARS-CoV-2, including pre- exposure of virus to samples for 1 hour in serum-free media prior to adding to cell plates. SARS-CoV-2 variants are tested in Vero E6 cells (ATCC) in MEM’s (Cytiva) supplemented with 50 μg/mL gentamicin (Sigma) and 2% FBS (Cytiva). Virus was diluted in serum-free test media to achieve a MOI of 0.001 for antiviral tests. Test peptides were serially diluted using eight 2-fold dilutions in serum-free test media. Serum-free was necessary to avoid peptide non-specific binding to components of FBS. Each dilution was added to 5 wells of a 96-well round-bottom plate containing no cells. Three wells of each dilution were infected with virus, and two wells remain uninfected as toxicity controls. Six wells were infected and untreated as virus controls, and six wells are uninfected and untreated as cell controls. Sample dilutions and virus are incubated together for 1 hour at 37C prior to adding to cell plates. Controls are tested in parallel. Following the pre-exposure of sample and virus, plate contents (100 μl per well) are transferred to plates containing 80-100% confluent Vero E6 cells with 100 μL MEM with 4% FBS and 50 μl/mL gentamicin (final culture media was MEM with 2% FBS and gentamicin on plate). Plates are incubated at 37C, 5% CO2. On day 4 post-infection, once untreated virus control wells reached maximum cytopathic effect (CPE), plates were stained with neutral red dye for approximately 2 hours. Supernatant dye was removed and wells rinsed with PBS, and the incorporated dye extracted in 50:50 Sorensen citrate buffer/ethanol for >30 minutes and the optical density was read on a spectrophotometer at 540 nm. Optical densities were converted to percent of cell controls and normalized to the virus control. The concentration of test compound required to inhibit CPE by 50% (EC50) was calculated by regression analysis. The concentration of compound that would cause 50% cell death in the absence of virus was similarly calculated (CC50). The therapeutic index (TI) was the CC50 divided by EC50.

### Molecular Dynamics

Explicit solvent molecular dynamics (MD) simulations of the novel coronavirus Spike protein were performed using the NAMD2 program^48^. We used the CHARMM-GUI ^49^ with the CHARMM36m force field along with TIP3P water molecules to explicitly solvate proteins and add any missing residues from the experimental structure files. Disulfide bonds and glycosylated sites were all included. Simulations are carried out maintaining the number of simulated particles, pressure, and temperature (the NPT ensemble) constant with the Langevin piston method specifically used to maintain a constant pressure of 1 atm. Periodic boundary conditions were employed and initial equilibration for a water box simulation. For water, the particle mesh Ewald (PME) method was used with a 20 A cutoff distance between the simulated protein and water box edge. The integration time step was 2 femtoseconds with our protein simulations conducted under physiological conditions (37 C, pH of 7.4, physiological ionic strength with NaCl ions, LYS and ARG were protonated and HIS was not). All mutations associated with Spike VOC’s and peptide residue substitutions associated with optimization were added via the CHARMM-GUI during the course of the pandemic.^49^ However, experimental Spike VOC structure files were used when available, such as Omicron PDB ID: 7T9L (Supplemental S2)^50^. The closeness of computationally generated structure versus experimental structure has also been recently demonstrated^51^. Our studies show that 100 nsec. was required to reach an equilibrium from mutated states from the WT structure files^52^. The total simulation time was therefore taken as 0.5 microseconds (250 million-time steps). All MD results given here were also repeated several times in order to help confirm trends in data.

### All-Atom Energetic Mappings-OpenContact

Previously,^52^ we analyzed the complete inter- and intra-protomer interactions across two independently published structure files (PDB ID: 6VSB and 6VYB; one up and two down protomers) for SARS-CoV-2 trimeric Spike protein using the open source energy mapping algorithm developed by Krall et al ^53^. This spatial and energetic mapping algorithm efficiently parses the strongest or most dominant non-covalent atom-atom interactions (charge and partial atomic charge, Born, and van der Waals forces), according to empirically established parsing criteria, based on the ab initio AMBER03 force field model. Following our previous studies, the parsing criteria was taken as the upper limit of -0.1 kT units for Lennard-Jones (van der Waals) criteria and -0.3 kT units for Coulombic interactions, although lower values can also be specified in the analysis part of the mappings in order to further refine the results.^53^ Note that in the all-atom analysis dominant van der Waals interaction forces are commonly associated with nonpolar atom-atom interactions and hydrophobic protein interaction regions, whereas the Coulombic partial charge and charge interactions are commonly associated with hydrophilic protein interaction regions and can include hydrogen bonding and backbone atom partial charge interactions.

## Supporting Information

S1. Mappings of human ACE2 to Spike RBD, SARS-CoV-2. (Please see Ref. 53 for more explanation on the data output.)

S2. Toxicity to A549 Cells and Computational Binding using Omicron PDB ID 7T9L

S3. Additional Supporting SPR data: hACE2 binding to Omicron RBD as positive control. S4. Raw Data: Authentic Virus Infectivity Assays.

## Acknowledgements

BSL3 studies reported here were carried out through ImQuest Biosciences (NJ). This work was supported, in part, by Hoth Therapeutics Inc., New York. Computations were carried out at the Center for High Performance Research Computing at VCU. Optimized peptides are associated with VCU Patent Applications: US 17/997,038 and PCT/US2021/024319. Special thanks to Mary Murphy (Reichart), and Mike Davis (VCU HPRC) for help and guidance.

## Author Contributions

MHP conceived and designed the peptides, performed experiments, and wrote the manuscript. XYP performed SPR experiments, data analysis, and helped review the manuscript.

## Notes

### Competing Interest Statement

The authors have declared no competing interest.

